# Conservation and Divergence of Related Neuronal Lineages in the *Drosophila* Central Brain

**DOI:** 10.1101/656017

**Authors:** Ying-Jou Lee, Ching-Po Yang, Yu-Fen Huang, Yisheng He, Qingzhong Ren, Hui-Min Chen, Rosa Linda Miyares, Hideo Otsuna, Yoshi Aso, Kei Ito, Tzumin Lee

**Affiliations:** Janelia Research Campus, Howard Hughes Medical Institute, Ashburn, VA 20147, USA

## Abstract

Wiring a complex brain requires enormous cell specificity. This specificity is laid out via a developmental process where neural stem cells produce countless diverse neurons. To help elucidate this process and resolve the considerable dynamic specificity, we need to observe the development of multiple neuronal lineages. *Drosophila* central brain lineages are predetermined, comprised of a fixed set of neurons born in pairs in a specific order. To reveal specific roles of lineage identity, Notch-dependent sister fate specification, and temporal patterning in morphological diversification, we mapped approximately one quarter of the *Drosophila* central brain lineages. While we found large aggregate differences, we also discovered similar patterns of morphological specification and diversification. Lineage identity plus Notch state govern primary neuronal trajectories, whereas temporal fates diversify terminal elaborations in target-specific manners. In addition, we identified ‘related’ lineages of analogous neuron types produced in similar temporal patterns. Two stem cells even yield identical series of dopaminergic neuron types, but with completely disparate sister neurons. These phenomena suggest that large changes in morphological diversity can be the consequence of relatively small differences in lineage fating. Taken together, this large-scale lineage mapping study reveals that relatively simple rules drive incredible neuronal complexity.

## Introduction

In order to understand how the brain encodes behavior, we need to study the developmental mechanisms that build and wire complex centers in the brain. The fruit fly is an ideal model system to research these mechanisms. The *Drosophila* field has extensive genetic tools; and a relatively small, yet complex fly brain enables scientists to connect neurons into functional circuits, map neural lineages, and test the role of essentially any gene in neurodevelopment and/or behavior [1]. Neurons are wired into neural networks that are connected through neuropils, neurite bundles with dense neurite arborization and numerous synapses. Neuropil structures in adult *Drosophila* brains have been extensively characterized, showing stereotypy and enabling alignment of multiple brains. Further, systematic efforts in cataloging neurons based on morphology and genetic drivers will soon yield a complete neuron type list for the fly central nervous system (CNS) [2]. Additionally, the fly brain connectome is being constructed at the level of individual synapses [3]. Finally, both morphology and development of the *Drosophila* brain are trackable at the single-cell level. It is therefore possible to resolve fly brain development from neural stem cells to the connectome and ultimately engineer the brain.

The *Drosophila* central brain develops from approximately 100 pairs of bilaterally symmetric neural stem cells, called neuroblasts (NBs) [4]. Most central brain NBs undergo around 100 cell cycles in two neurogenic periods, the first to build larval networks and then more complex adult neural networks [5]. A typical neuronal lineage, born from a single NB, consists of serially derived pairs of post-mitotic neurons. Due to differential Notch signaling, the paired neurons, made by transient ganglion mother cells (GMCs), can acquire very distinct morphological, physiological, and molecular phenotypes depending on lineage-specific binary sister fates, referred to A (N^on^) versus B (N^off^) fate [6]. Neurons of the same A or B fate constitute a hemilineage [7]. Temporal patterning allows further neuronal fate diversification within a given hemilineage based on neuronal birth order [8]. Such lineage-dependent fate specification and diversification underlie the construction of the fly brain connectome.

Brain neurogenesis is spatiotemporally patterned and involves similar neural developmental genes across evolution. Fate-mapping distinct neural stem cell pools as well as birth-dating serially derived neurons also show lineage-guided neural fate specification and diversification in vertebrate brains [9]. However, it remains extremely challenging to track vertebrate brain development with single-cell resolution.

It is believed that lineages make up individual functional units in *Drosophila*. This was made obvious by marking clonally related neurons produced from individual NBs. NB clones are stereotyped with characteristic cell body positions and distinct lineage-specific neurite projections. Using MARCM [10], we identified approximately 100 distinct, full-size NB clones per central brain [11]. This indicates that each of the ~200 NBs consistently yields lineage-specific progeny. Such lineage identities arise from the spatially patterned neuroectoderm and involve evolutionarily conserved anteroposterior gap genes and dorsoventral patterning genes [12] [13]. Individual NBs in early embryos can indeed be identified based on combinatorial expression of various transcription factors (TF) [4]. Altering the TF code can transform neuronal fate and wiring, suggesting that lineage identity controls aspects of neuronal type and morphology [14]. Interestingly, fly and beetle have conserved NBs but express different TFs, further suggesting that NBs and the derived lineages can readily evolve with minor changes in gene expression [15].

To reveal how NB lineages guide neuronal diversification further requires mapping single neurons and their developmental origins. Labeling GMC offspring as isolated single-neuron clones and assigning them to specific lineages based on lineage-characteristic morphology are possible but tedious, and could be challenging for closely related lineages. Targeted cell lineage analysis using lineage-restricted genetic drivers is therefore preferred for mapping specific neuronal lineages of interest with single-cell resolution [16]. To date, only three of the about 100 distinct neuronal lineages have been fully mapped at the single-cell level in adult fly brains, the mushroom body (MB), anterodorsal antennal lobe (ALad1) and lateral antennal lobe (ALl1). These mapped lineages yield 1) MB Kenyon cells (KC), 2) AL projection neurons (PN), and 3) AL/AMMC PNs and AL local interneurons (LN), respectively. All three lineages sequentially produce morphologically distinct neuron types, indicating a common temporal cell fating mechanism. However, the progeny’s morphological diversity varies greatly from one to another lineage. The four identical MB lineages are composed of only three major KC types and paired KCs from common GMCs show no evidence for binary sister fate determination [17]. By contrast, the two AL NBs produce progeny that rapidly change type (producing upwards of 40 neuron types) and the GMCs generate discrete sister fates [17]. In the ALl1 lineage, differential Notch signaling specifies PNs versus LNs in the ALl1 lineage [18] [19]. Notably, the paired PN and LN hemilineages show differential temporal fate patterning. Moreover, the ALl1 PN hemilineage consists of alternating AL and AMMC PNs, as a result of Notch signaling [19]. Together, these phenomena demonstrate a great versatility in lineage-guided neuronal diversification.

Assembling complex region-specific intricate neural networks for an entire brain requires enormous cell specificity. In fact, the cellular diversity as characterized by gene expression is higher in developing than mature brains [20], signifying that the underpinnings of the connectome can be understood by studying development. Such developmental diversity is reflected by characteristic neurite projection and elaboration patterns. We therefore aim to elucidate the roles of NB lineage specification, temporal patterning and binary sister fate decisions upon neuronal morphology. By doing this, we hope to gain insight about how a limited number of NBs can specify such enormous brain complexity. We chose to map a large subset of NB lineages, enough to make generalizations but not so many to confound analysis. To this end, we selected NBs expressing the conserved spatial patterning gene *vnd* [4]. With single-neuron resolution, we mapped 25 hemilineages derived from 18 vnd-expressing NBs. We observed hemilineage-dependent morphological diversity at two levels. First, neurons of the same hemilineage may uniformly innervate a common neuropil or differentially target distinct neuropils. Second, neurons show further structural diversity in terminal elaboration, which depends on neuropil targets rather than lineage origins. Once you factor in the differences of the neuropil targets, grossly distinct hemilineages show comparable patterns in the diversification of neuron morphology. Moreover, we discovered pairs of non-sister hemilineages that make similar or even identical neuron types with common temporal patterns. These observations suggest involvement of conserved lineage-intrinsic cell fating mechanisms in the derivation of diverse neuronal lineages for making complex brains.

## Results

### Mapping 18 neuronal lineages concurrently with *vnd-GAL4*

In order to target a large subset of related neuronal lineages, we wanted to exploit a conserved patterning gene. Both anteroposterior and dorsoventral patterning of the CNS are remarkably conserved from insects to humans [12] [13], including the tripartite organization of the brain (the forebrain, midbrain and hindbrain correspond to the fly’s protocerebral, deutocerebral, and tritocerebral neuromeres). With the aim of making our findings applicable to all three neuromeres, we searched for a conserved dorsoventral patterning gene with relatively even distribution. Urbach and Technau reported 21 fly brain NBs expressing Ventral nervous system defective (Vnd, homolog of the Nkx2 family of homeobox transcription factors). These include 13 of the 72 protocerebral NBs, 4 of the 21 deutocerebral NBs, and 4 of the 13 tritocerebral NBs [4]. Therefore, we decided to target these Vnd expressing NB lineages for a detailed, large scale analysis.

To analyze Vnd^+^ NBs, we created a GAL4 under the control of endogenous *vnd* regulatory sequences via gene targeting [21]. To immortalize the NB expression of Vnd into the neuronal progeny, we derived a Vnd-specific, lineage-restricted LexA driver using a cascade of site-specific recombinases. This cascade is triggered by vnd-GAL4, filtered through dpnEE (a pan-NB promoter), and then driven ubiquitously so that each of the NB’s daughter cells express LexA (Figure 1A). To isolate/identify individual Vnd lineages, we utilized stochastic clonal induction of a conditional LexA reporter. We detected 18 stereotyped neuronal lineages with cell bodies within the *Drosophila* central brain (data not shown). These lineages correspond to the SMPad1, SMPp&v1, SLPpm3, CREa1, CREa2, WEDd1, AOTUv1, AOTUv3, AOTUv4, VLPa2, VESa1, VESa2, ALv1, LALv1, FLAa1, FLAa2, FLAa3, and WEDa1 lineages that we previously identified [11]. The cell body clusters of these vnd-GAL4 lineages cover the medial part of the anterior brain surface, consistent with Vnd’s expression around the midline of the embryonic CNS.

**Figure 1.**
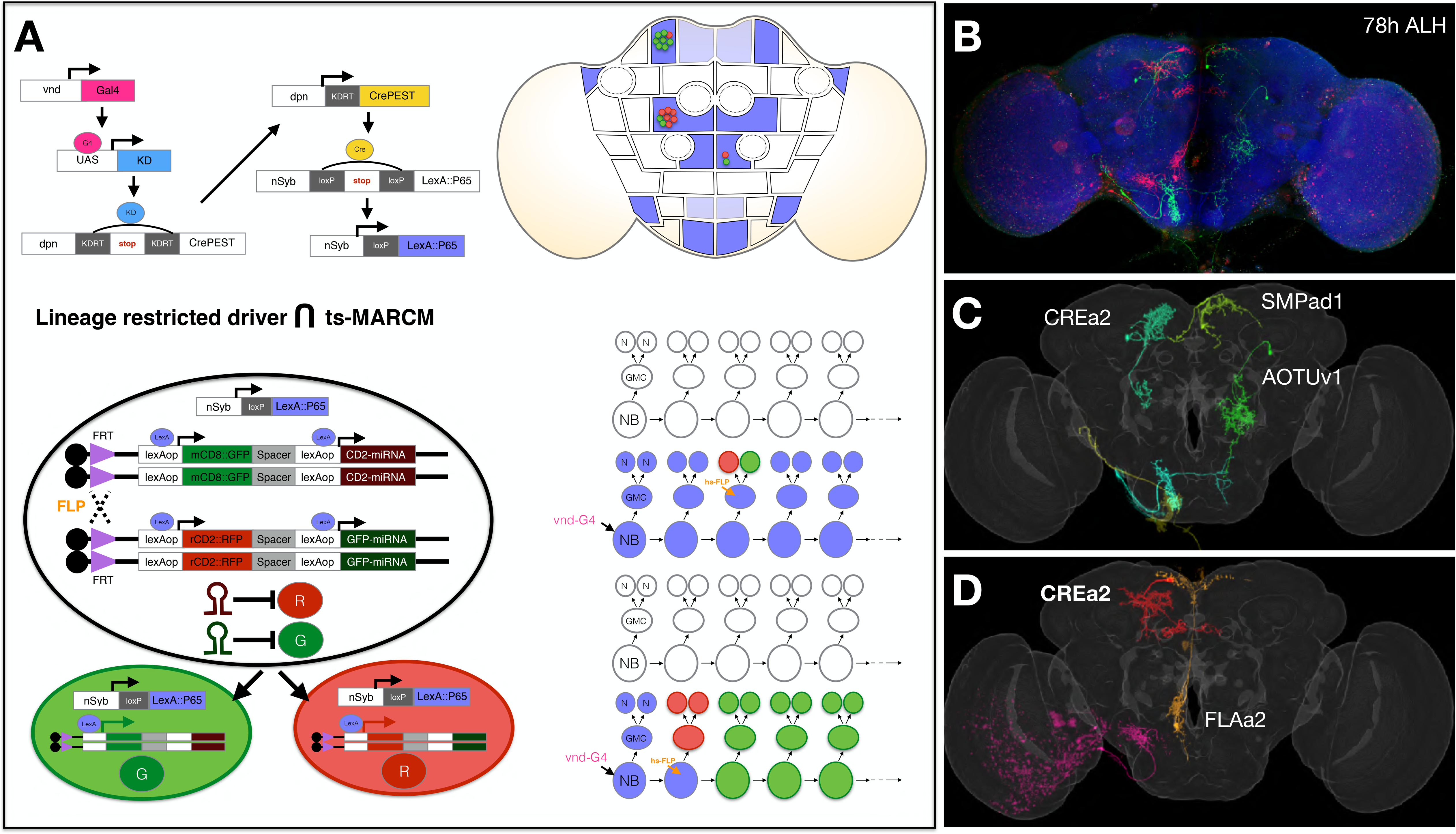
Concurrent mapping of Vnd neuronal lineages with twin-spot MARCM. (A) Schematic illustration of twin-spot MARCM with vnd-specific lineage-restricted driver. Left top: vnd-GAL4 reconstitutes dpn-CrePEST that in turn reconstitutes nSyp-LexA::P65. LexA::P65 is only expressed in neurons produced by Vnd^+^ NBs derived from vnd-expressing domain. Left bottom: mitotic recombination leads to differential labeling of paired sister cells by twin-spot MARCM. Right top: only in those lineages positive for nSyb-LexA::P65 (blue intensity indicating frequency of nSyb-LexA::P65 reconstitution) can the stochastically induced, paired homozygous clones be differentially labeled as twin spots. Right bottom: mitotic recombination in GMC vs. NB elicits paired single-cell clones vs. GMC offspring (red) paired with the remaining lineage (green). (B) A representative nc82-counterstained (blue) adult fly brain carrying multiple twin-spot clones (green/red), induced at 78h ALH. (C) Various green neurons from [B] were segmented out and then warped into a standard adult fly brain. We annotate lineage origin for neurons with cell bodies in the central brain. (D) Various red neurons from [B] were segmented out and then warped into a standard adult fly brain. We annotate lineage origin for neurons with cell bodies in the central brain.

Despite sharing Vnd expression, the labeled NB clones each show distinct gross morphology. To unravel the extent to which lineage origins and temporal regulation govern neuronal morphology and target selection, we compared the progeny of each Vnd^+^ NB in detail. We identified individual neurons based on morphology and determined the birth order for each of the 18 vnd-GAL4 lineages. We simultaneously mapped all Vnd lineages by twin-spot MARCM using the Vnd-specific, lineage-restricted LexA driver (Figure 1). We conducted transient clone induction in contiguous two-hour windows from 18h after larval hatching (ALH) to 92h ALH and from 22h before pupa formation (BPF) to 16h after pupa formation (APF). We imaged 5771 brains containing twin-spot clones of the Vnd 18 lineages. We manually annotated individual clones (e.g. Figure 1B-D) and ascribed 20,916 clones to specific Vnd lineages in the brain. Although the total clone number for a given lineage varied from hundreds to thousands, these differences likely resulted from differential apoptosis of the progeny or different lineage length. For instance, the VESa1 and VESa2 clones were almost exclusively induced prior to 80h ALH, implicating stage-specific progeny production or viability. These observations indicate a rather homogenous coverage of the 18 brain lineages with *vnd-GAL4*.

Twin-spot MARCM utilizes mitotic recombination to independently label the progeny of a cell division in different color [22]. This can occur in a cycling NB (labeling the progeny of a GMC in one color and the remainder of the lineage in another) or it can occur in a GMC (labeling the two daughter neurons in different colors). However, programmed cell death could result in unpaired or single-neuron-paired NB clones or isolated single-cell clones. In fact, only seven of the 18 Vnd lineages (SMPp&v1, CREa1, CREa2, AOTUv1, AOTUv3, AOTUv4, and LALv1) produce two viable hemilineages. Regardless of lineage origin, the paired neurons across sister Vnd hemilineages consistently exhibit distinct morphology and have different neuropil targets, reflecting Notch-dependent binary sister fates. The remaining 11 lineages exist as a lone hemilineage, as evidenced by the recovery of only unpaired single-neuron clones sharing lineage-specific primary trajectories and often innervating common neuropils. Given the drastic hemilineage distinctions, our analyses of neuron diversity considered the 25 hemilineages separately. Nonetheless, in the seven lineages composed of sister hemilineages, we exploited the paired sister neurons to determine lineage identity and to compare sister hemilineage development.

We further addressed the hemilineages’ Notch state by silencing Notch with RNAi in isolated NB clones. Repressing Notch should promote Notch^off^ fate, resulting in a reduction/loss of the Notch^on^ morphology and increasing the number of neurons with Notch^off^ morphology. However, when examining NB clones consisting of two sister hemilineages, we noticed that a severe reduction in one hemilineage’s morphology might not be accompanied by an evident enhancement of the other. This could be due to death of ectopic neurons because of an incomplete transformation or limited space/resources. Further, the reduction in the number of neurons with N^on^ morphology was incomplete, likely due to weak RNAi. This was particularly obvious among neurons born shortly after clone induction. Nonetheless, we could confidently resolve the Notch state for most of the 25 Vnd hemilineages, including all the 14 hemilineages that exist in pairs. For simplicity, we pseudo-colored neurons derived from Notch^on^ hemilineages in green, neurons from Notch^off^ hemilineages in red, and neurons from uncertain hemilineages in yellow.

### Morphological complexity decreases with birth order

To map progeny diversity, we clustered single-neuron clones based on neuron morphology and timing of clone induction. For each of the 18 Vnd lineages, we identified morphologically distinguishable neuron types and further determined their approximate birth sequence based on the peak production time of each neuron type. Single neurons consistently occupied a much more restricted domain compared to full-size NB clones. Moreover, single-neuron clones exhibited birth order-dependent trajectories. Given these phenomena, we examined the extents of lineage coverage in our single-neuron collection by merging representative single neurons from all annotated morphological types and comparing the merged single cells with full-size NB clones. Aligning samples through a standard 3D fly brain template confirmed that all major trajectories from a NB clone were covered by single neuron projections, ensuring we have not missed any major neuron types. Together, our analysis demonstrates that we have simultaneously mapped the 18 Vnd lineages with single-cell resolution.

In the process of overlaying single-neuron clones onto full-size NB clones, we noticed that the first larval-born neurons show uniquely distant elaborations in eight (32%) hemilineages, including SLPpm3, CREa1(N^off^), ALv1, VESa1, VESa2, FLAa1, FLAa2, and VLPa2. These first larval-born neurons consistently extend farther than the remaining offspring. We thus see extra distant targets exclusively on the GMC side of twin-spot NB clones induced around quiescence exit (Figure 2). In another five hemilineages including CREa1(N^on^), CREa2(N^off^), AOTUv1(N^on^), AOTUv4(N^on^), and WEDd1, the above “first-born” phenomenon extends longer, over several early-born neurons. For instance, the postembryonic WEDd1 lineage consistently start with two descending neurons with ventral nerve code (VNC) projections, followed by neurons completely confined to the brain. All together, we found that 13 out of 25 (52%) Vnd hemilineages contain neurons with uniquely exuberant elaborations born at the beginning of larval neurogenesis.

**Figure 2.**
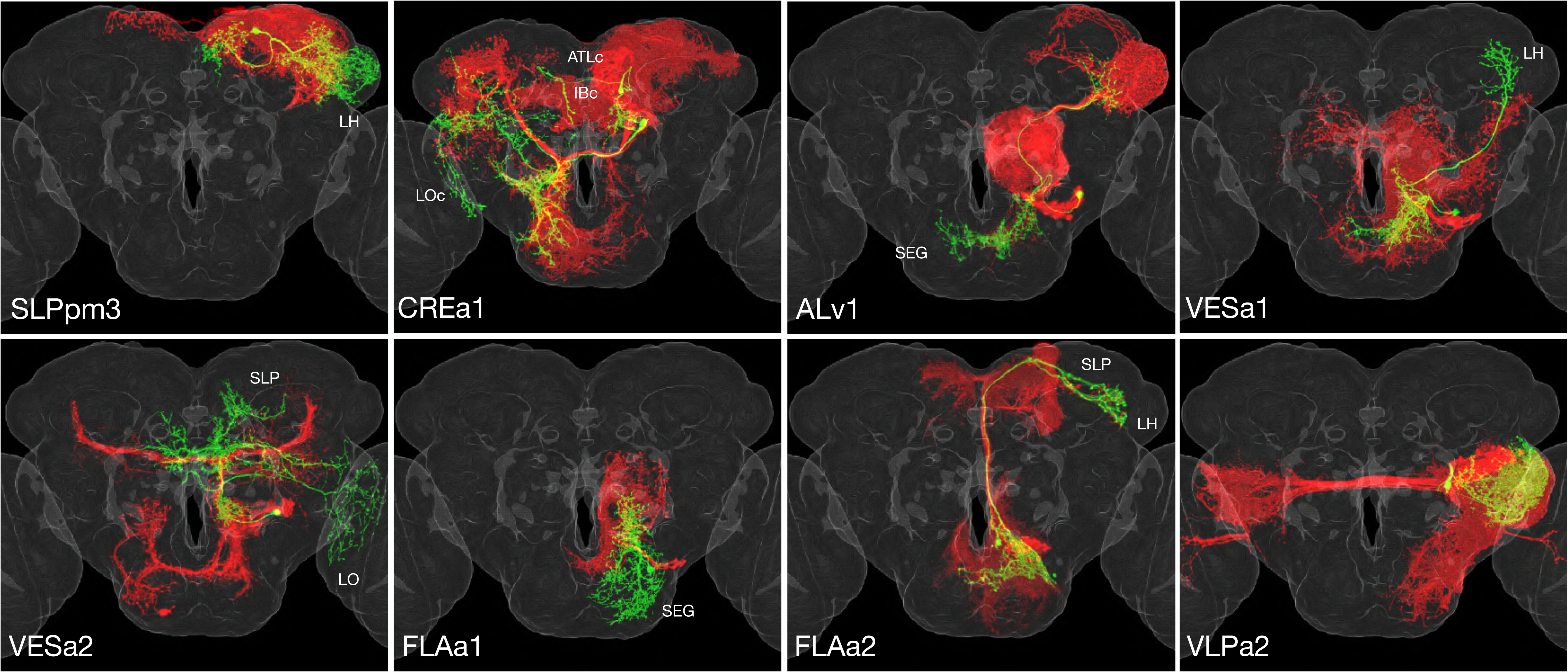
Uniquely exuberant first larval-born neurons. Isolated twin-spot clones shown in adult fly brain template. Each clone consists of one or two (only CREa1) first larval-born neurons (green) paired with all subsequently born neurons (red) of the same lineage. Neuropils uniquely innervated by the shown first larval-born neurons are indicated.

Contrasting the complexity of early-born neurons, seven hemilineages (28%), including SMPad1, SLPpm3, CREa1(N^off^), CREa2(N^off^), AOTUv1(N^off^), LALv1(N^off^), and VESa1, terminate with neurons that have severely reduced domains of elaboration. Three of them, including SLPpm3, CREa1(N^off^), and VESa1 (Figure 3), have produced a uniquely exuberant first larval-born neurons, then many neurons with intermediate elaboration, and lastly neurons with limited elaboration. Taken together, our data suggests that the extent of neuronal elaboration is regulated by temporal fate in a similar manner across diverse lineages.

**Figure 3.**
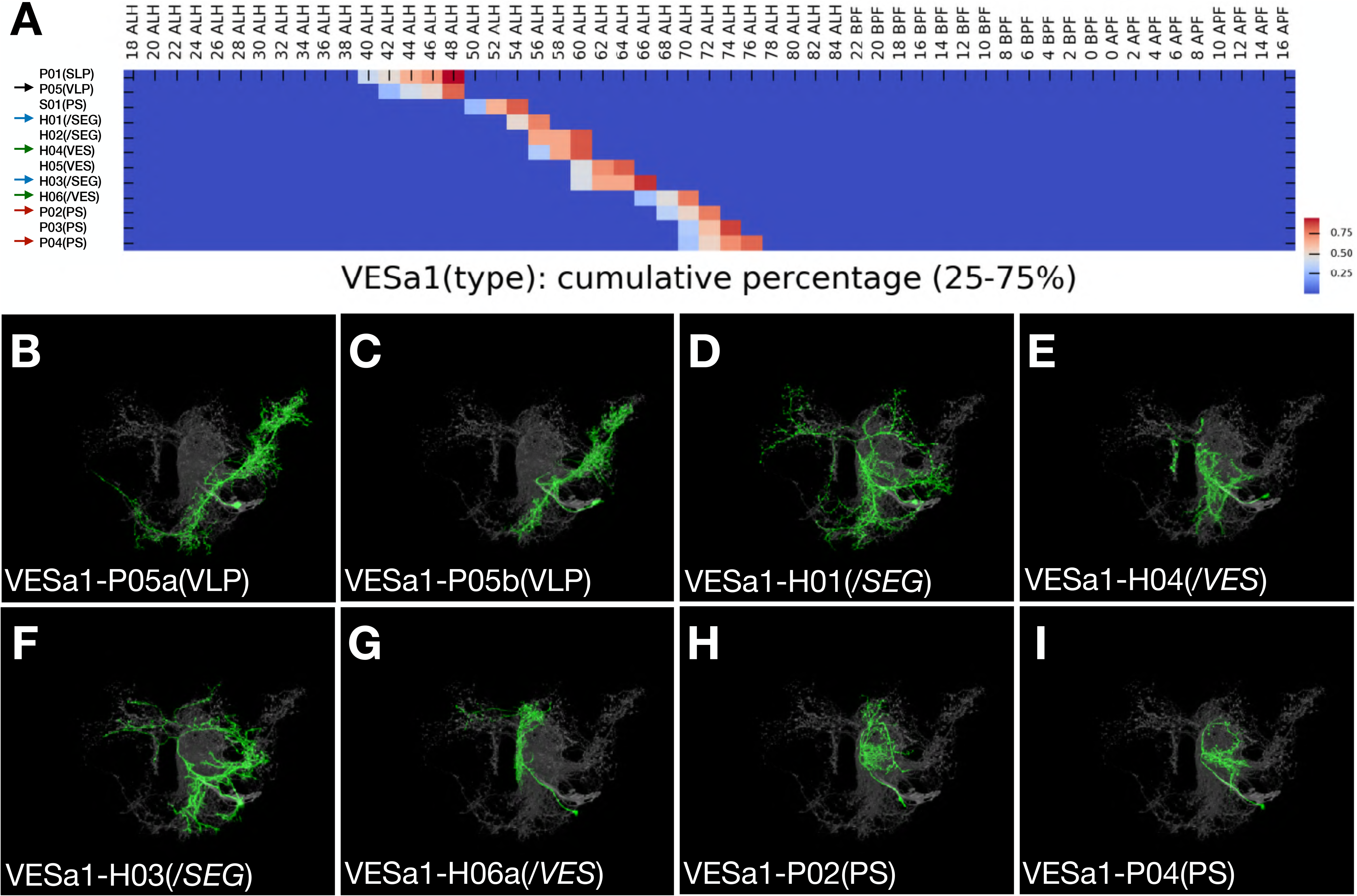
Late-born neurons show simplified morphology. (A) Heatmap showing birth time distribution of specific VESa1 neuron types with cumulative coverage of 25% or higher (until passing 75%). Arrows indicate neuron types shown in [B-I]. See text for neuron type nomenclature. (B-I) Serially derived single neuron types (green) shown in the context of all recovered VESa1 neuron types (grey). Note reduction in neurite elaboration along birth time.

### Neurons of same hemilineage origin vary in topology

To what extent do lineage origins determine neuronal morphology and neuronal targets? Considering neuron topology in the context of brain-wide networking, we established a refined neuron classification scheme that takes into account neuron topology and basic topographic features (Figure 4). First, we assign those brain-input/output ‘extrinsic’ neurons, including descending neurons, to the External cluster. Second, we classify brain-intrinsic neurons based on (1) single or multiple domains of arborization, and (2) unilateral, bilateral, or midline targeting. Briefly, we designate neurons with a single unilateral arborization domain as Single (S), neurons with multiple arborization domains exclusively within one hemisphere as Projection (P), neurons with a single domain of arborization covering the brain midline as Central (C), neurons with midline targeting plus non-midline arborization as Midline (M), and neurons with midline crossing as either Transverse (T) or Horizontal (H) depending on absence or presence of bilaterally symmetric innervation.

**Figure 4.**
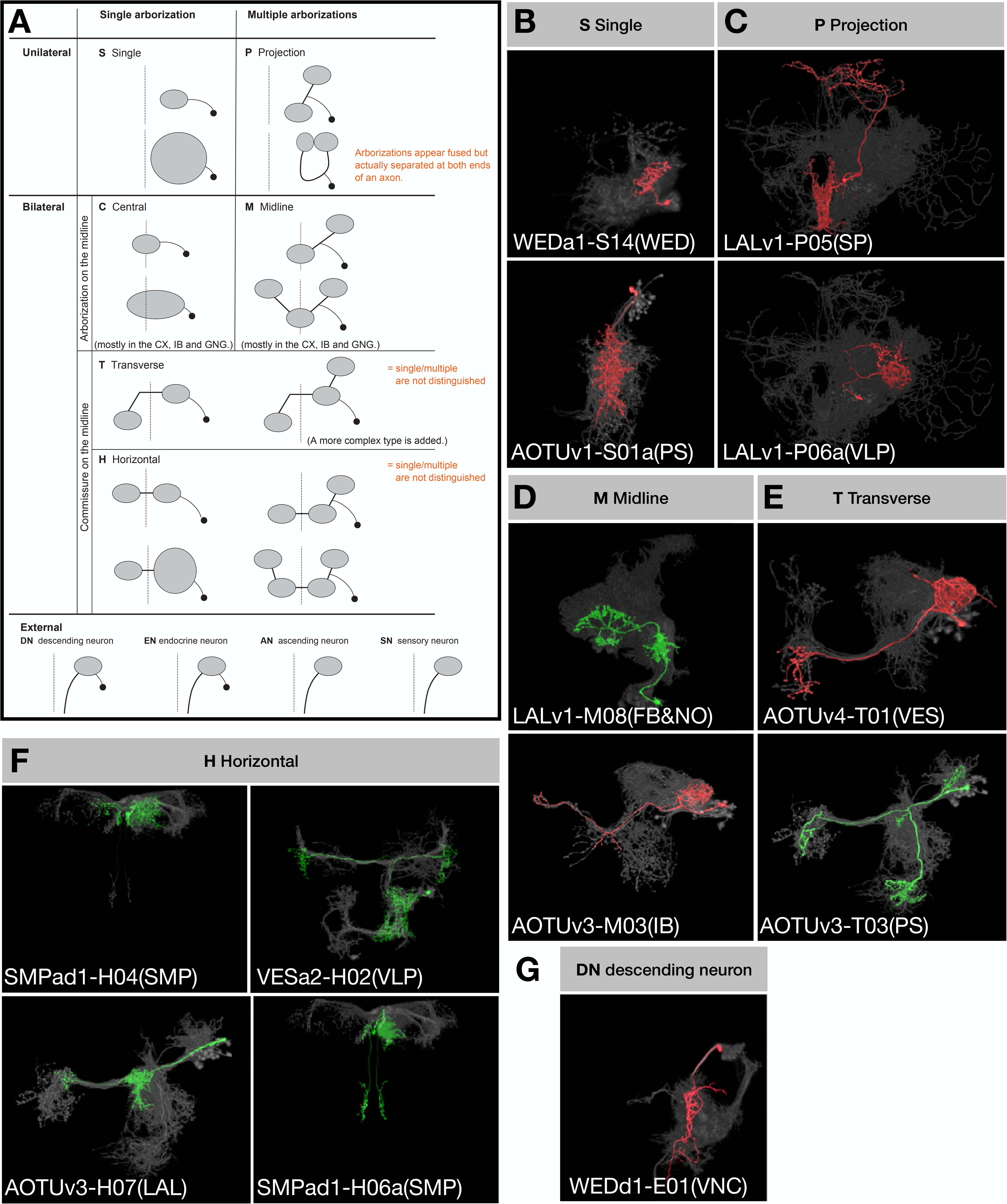
Topological classification of single neurons. (A) Schematic illustration of neuron morphology classification. Dashed lines indicate the brain midline. Black dots: cell bodies; grey ovals: neurite elaborations. (B-G) Representative single neuron types (green: N^on^, red: N^off^) shown in the context of all recovered neurons (grey) of the same hemilineages (indicated in the beginning of neuron type names). One neuron type corresponding to each illustrated class is shown. Note no C-topology neurons were found in the 18 vnd lineages, and for External neurons, only the DN type was found.

We annotated the neurons of Vnd lineages according to this classification scheme. We found numerous P-class, H-class, and M-class neurons, many T-class neurons, a few S-class neurons, and only two descending neurons. Notably, only five (20%) Vnd hemilineages consist of neurons that exclusively belong to a single topological class. If we exclude the extremely elaborate first-born and austere last-born neurons, we can add four additional hemilineages into this list of topologically pure hemilineages. By contrast, there are 11 (44%) Vnd hemilineages of three to four topological classes of neurons on top of those extreme beginning and ending neurons. Taking the SMPp&v1 sister hemilineages as examples, we see P, H and S classes of N^on^ progeny but T, H and M classes of N^off^ progeny (Figure 5). Comparing sister hemilineages, the H-class N^on^ and N^off^ cousin neurons display no morphological resemblance despite belonging to the same topological class. By contrast, within a given hemilineage, neurons of various classes often have strong resemblances. For instance, some H-class and P-class SMPp&v1(N^on^) neurons have nearly indistinguishable elaborations on the ipsilateral side. Common ipsilateral morphology also exists among the H, T and M classes of SMPp&v1(N^off^) neurons. Moreover, the S-class neurons look like simplified P-class neurons within the SMPp&v1(N^on^) hemilineage. These phenomena demonstrate the versatility of neurons in adopting various topologies, thus allowing frequent coexistence of H/T/M, P/H, P/S, or H/S classes.

**Figure 5.**
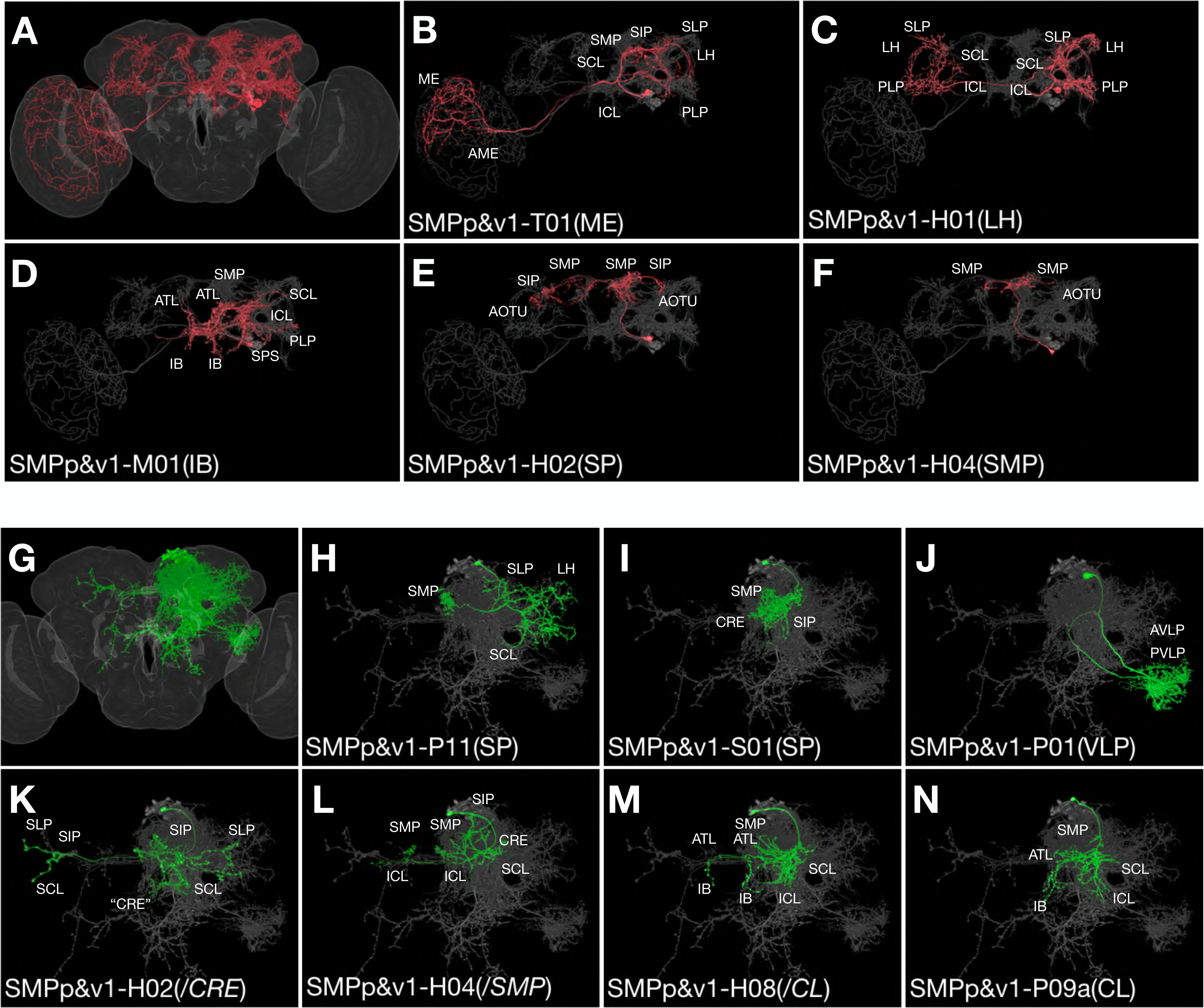
Clonally related neurons show diverse topologies. (A) All recovered SMPp&v1B neuron types merged onto the standard fly brain template. (B-F) Serially derived single neuron types (red) shown in the context of all recovered SMPp&v1B neuron types merged together (grey). Note presence of T-, M-, H-topolgy neurons with comparable ipsilateral elaborations. (G) All recovered SMPp&v1A neuron types merged onto the standard fly brain template. (H-N) Serially derived single neuron types (green) shown in the context of all recovered SMPp&v1A neuron types merged together (grey). Note presence of P-, S-, H-topology neurons in the same hemilineage, and similarities among certain S-, H-, and P-topology neurons (I,M,N).

Given the hemilineage-dependent coexistence of selective topological classes, we decided to name clonally related morphological types of neurons with the following convention. Our nomenclature starts with lineage name followed by topology-class letter and a two-digit number. The lineage name is separated from the topology-class letter with a dash sign. To distinguish sister hemilineages, we add ‘A’ or ‘B’ after the lineage name to represent N^on^ or N^off^. The two-digit serial number starts from 01 and is assigned by arranging neuron types of the same hemilineage-topology class through first clustering neuron types based on closeness in morphology and then determining cluster sequence as well as intra-cluster neuron sequence to roughly reflect birth order. We typically provide extra information in parentheses at the end, to indicate the key morphological feature (e.g. the most distal neuropil target) or simply the preexisting name if available. Briefly, we have identified 467 morphological neuron types from 25 vnd hemilineages in the fly central brain (supplemental table). The five most heterogeneous hemilineages include: ALv1 (46 types), VLPa2 (37 types), LALv1B (31 types), LALv1A (29 types), and CREa1A (26 types). The five least heterogeneous include: FLAa1 (5 types), FLAa3 (7 types), CREa1B (9 types), SLPpm3 (11 types), and VESa1 (12 types). The remaining 15 hemilineages yield 15 to 24 morphological neuron types. However, we could have over-estimated neuron types due to structural plasticity or under-estimated neuron types due to lack of landmarks, especially in neuropils that are poorly characterized.

### Hemilineages vary greatly in gross complexity

A ‘diverse’ hemilineage might consist of distinct neurons that uniformly target the same set of neuropils or grossly dissimilar neurons each innervating distinctive sets of neuropils. Given this phenomenon, we further cluster clonally related neuron types into morphological groups based on patterns of neuropil targeting. For instance, the paired LALv1A and LALv1B hemilineages yield similar numbers (29 vs. 31) of morphologically distinguishable neuron types. However, the LALv1A hemilineage produces only one dominant morphological group, whereas LALv1B neurons can be clustered into eight morphological groups, including five large groups plus three small groups.

Notably, all LALv1A neurons born after larval hatching project along the same primary track into the central complex (CX), including fan-shape body (FB), noduli (NO), ellipsoid body (EB), and asymmetric body (AB) (Figure 6A-L). The majority of LALv1A neuron types innervate specific FB layers. These FB neurons are preceded in birth order by several NO- or EB-targeting neurons, and followed by multiple AB-targeting neurons. Within the large middle FB window, there are 26 distinguishable neuron types that arise in an invariant sequence. The first three types innervate FB layer 3/4 and ipsilateral NO and show distinguishable proximal elaborations. The fourth neuron type is an outliner, skipping the FB and uniquely targeting the contralateral NO. The next six types selectively innervate a single FB layer, serially targeting layers 2, 4, 7, 6, 5, and 8. The following four types display two-layer targeting, and the next four types selectively target layer 3 or 4. A similar sequence with different multi-layer FB neurons followed by single-layer FB neurons (targeting layer 2 or 1) repeats once. Interestingly, FB layer 4 receives innervation from nine LALv1A neuron types. By contrast, most FB layers are innervated by only three types. Further, layer 4 receives major inputs from CRE/LAL/VES whereas other FB layers receive inputs from SP/CRE/LAL. In sum, all FB layers are innervated by two-layer as well as single-layer FB neurons arising from LALv1A in a specific sequence, yet with unclear logic.

**Figure 6.**
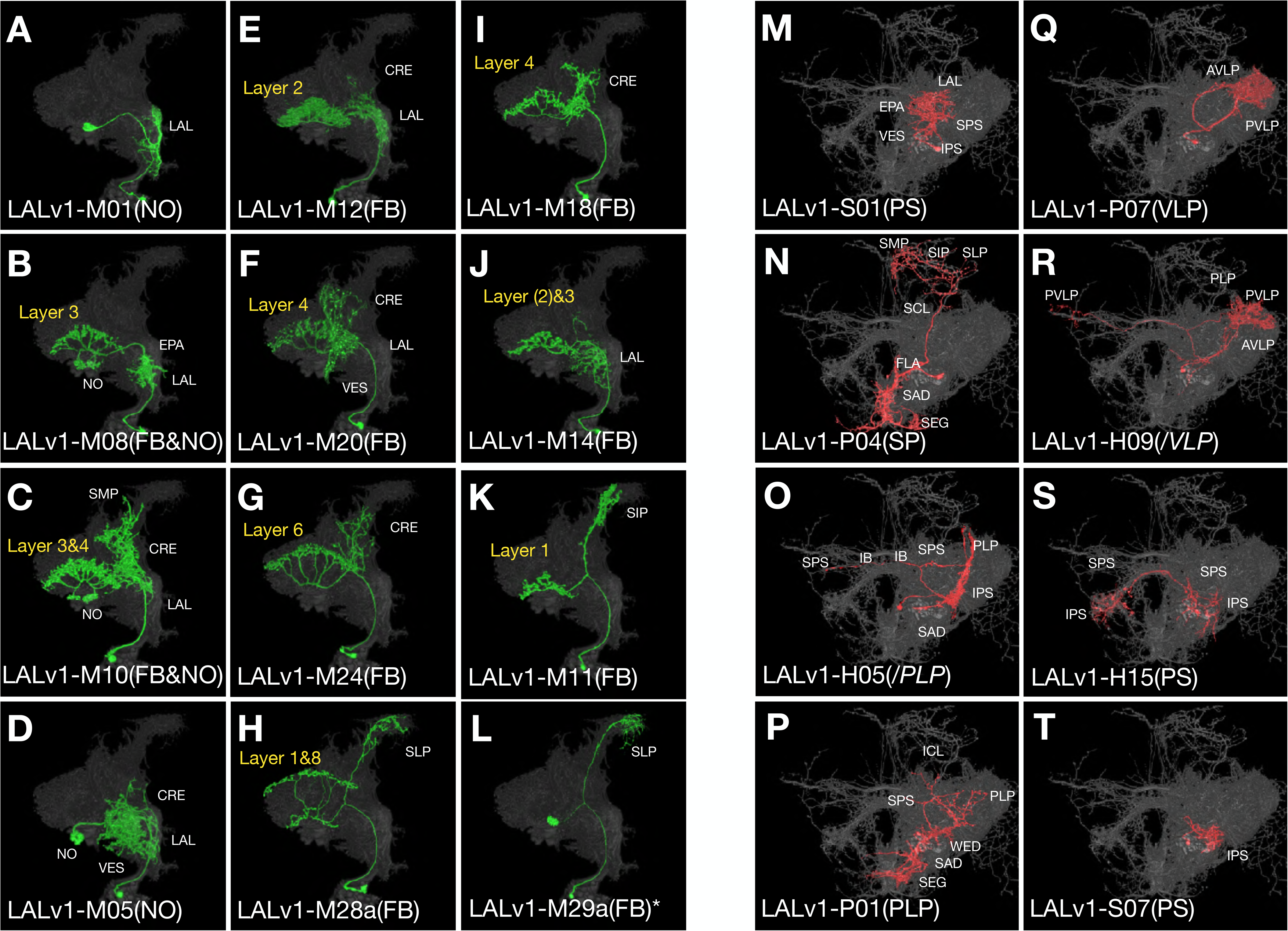
Uniform CX neurons pair with diverse groups of non-CX neurons in LALv1. (A-L) Serially derived single neuron types (green) shown in the context of all recovered LALv1A neuron types merged together (grey). Note serial innervation of specific CX subcompartments, recurrent targeting of some subcompartments (e.g. layer 4 of the FB in [F,I]), and innervation of the asymmetric body (AB) by the last-born type (L). (M-T) Serially derived single neuron types (red) shown in the context of all recovered LALv1B neuron types merged together (grey). Note orderly production of the five morphological groups that respectively target PS, SP, PLP, VLP, and PS again. Neurons of the same morphological group often show shared morphological features (e.g. [Q] vs. [R] and [S] vs. [T]).

In contrast to the relatively uniform LALv1A hemilineage, the sister hemilineage, LALv1B, consists of five major morphological groups with distinct patterns of neuropil targeting (referred to as PS1, SP, PLP, VLP, and PS2 group based on the main neuropil of innervation) (Figure 6M-T). These five major groups arise sequentially following the birth of four neuron types with three additional patterns of neuropil targeting. Neurons in a given morphological group can vary in topology, implicating again the versatility in adopting related topologies. However, different morphological groups contain distinct group-characteristic topological classes. This point is reaffirmed by the differential presence of P versus H class in the two temporally separated PS groups. Further, we see similar-looking S-class neurons in both PS groups, which could however arise through a different topological reduction from P versus H to S. Nonetheless, the multiple morphological groups of related neuron types in the LALv1B hemilineage arise sequentially in a progressive manner.

Cross comparison of LALv1A and LALv1B revealed multiple windows of uncoordinated temporal fate changes across sister hemilineages. There exist windows with a series of distinct LALv1A neurons paired with LALv1B neurons of the ‘same’ type, and vice versa. Given the well-defined stereotyped organization of CX sub-compartments, we have high confidence that we identified all individual LALv1A neuron types based on morphology. However, we may have under- or over-estimated the number of LALv1B neuron types. For example, there could exist unidentified types of VLP-targeting LALv1B neurons with the P topology, as we only distinguished two types of P-class VLP neurons from the same window of time where the sister LALv1A hemilineage contains seven types of FB neurons. Conversely, we might have overestimated the number of PS-targeting LALv1B neuron types, especially among late-born neurons which appeared more plastic in morphology. Nonetheless, at the level of morphological groups, the paired LALv1A and LALv1B hemilineages show independent temporal patterning despite their derivation form a common neural stem cell.

### 12 of 25 hemilineages yield only one dominant morphological group

There are 11 additional (48% in total) Vnd hemilineages which, like LALv1A, contain only one dominant morphological group. Notably, four of the above five most heterogeneous Vnd hemilineages (having the highest number of distinguishable neuron types), including ALv1, VLPa2, LALv1A, and CREa1A, are among the 12 hemilineages with rather uniform single-neuron morphology. This paradoxical phenomenon evidently results from the composition of many neuron types involved in constructing fine topographic maps. In particular, the postembryonic VLPa2 hemilineage is exclusively dedicated to the formation of the visual topographic map in the VLP neuropil (Figure 7). Analogously, 45 of the 46 ALv1 neuron types (except the unique first larval-born neuron) uniformly relay olfactory information from the AL to LH.

**Figure 7.**
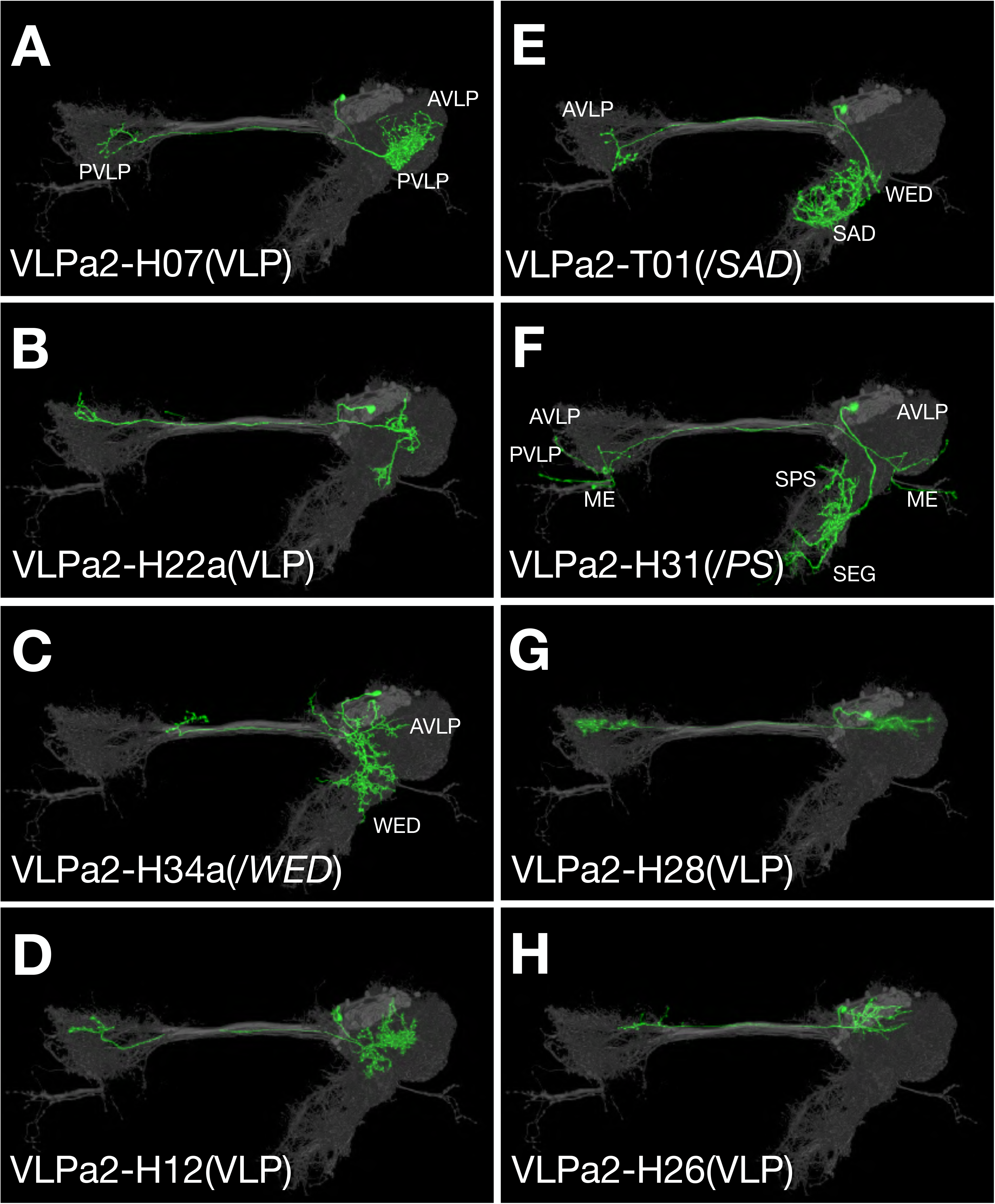
Typical VLPa2 neurons show symmetric targets with more extensive ipsilateral elaboration. (A-H) Representative VLPa2 neuron types arranged based on birth order. Only one (E) out of the shown eight types adopts T-topology and lacks bilateral VLP innervations.

Besides VLP and LH, the main neuropils targeted by relatively uniform Vnd hemilineages include: the FB (innervated by LALv1A and AOTUv4A), MB lobes (innervated by CREa1A and CREa2A), SP (innervated by SMPad1, SLPpm3, FLAa3, and CREa1B), PS (innervated by AOTUv1), and VES (innervated by the small FLAa1 hemilineage). Below, we present AOTUv4 and describe the co-innervation of MB lobes by the CREa1A and CREa2A hemilineages. As to innervation of SP, PS, and VES, we are uncertain of the significance of having uniform hemilineages because little is known about these neuropils.

### 13 of 25 hemilineages yield multiple neuron groups targeting discrete neuropils

There are 12 additional (52% in total) Vnd hemilineages which, like LALv1B, contain multiple neuron groups targeting discrete sets of neuropils. This list includes five unpaired hemilineages (VESa1, VESa2, FLAa2, WEDd1, WEDa1), four paired with a uniform hemilineage (LALv1B, CREa2B, AOTUv1A, AOTUv4B), and two pairs of sister hemilineages (SMPp&v1, AOTUv3). Examining the temporal patterns of neuron morphological diversity revealed progressive neuropil targeting in 11 hemilineages and recurrent neuropil targeting (same pattern in multiple windows) only in AOTUv3A and VESa2. Below, we utilize the paired AOTUv3B and AOTUv3A hemilineages to illustrate these two distinct patterns of birth order-dependent neuropil targeting.

The AOTUv3B hemilineage encompasses six sequential morphological groups of neurons (Figure 8A-H). The first four groups almost exclusively consist of P-topology neurons, which relay information from AOTU or SP to LAL or from AOTU to CRE or BU. However, there is a lone M-topology neuron type with a SP-to-IB/CRE/LAL projection pattern born at the transition from the SP-to-LAL P-topology group to the AOTU-to-CRE P-topology group. Nonetheless, this ‘ectopic’ M-topology neuron does resemble the preceding SP-to-LAL P-topology neurons in other aspects, reminiscent of versatile neuron topology. The following two groups only contain M-topology neurons, which connect AOTU with IB/SP and SP with IB/ATL respectively. Taken together, the AOTUv3B hemilineage displays clean, progressive transitions to produce neurons with diverse morphology.

**Figure 8.**
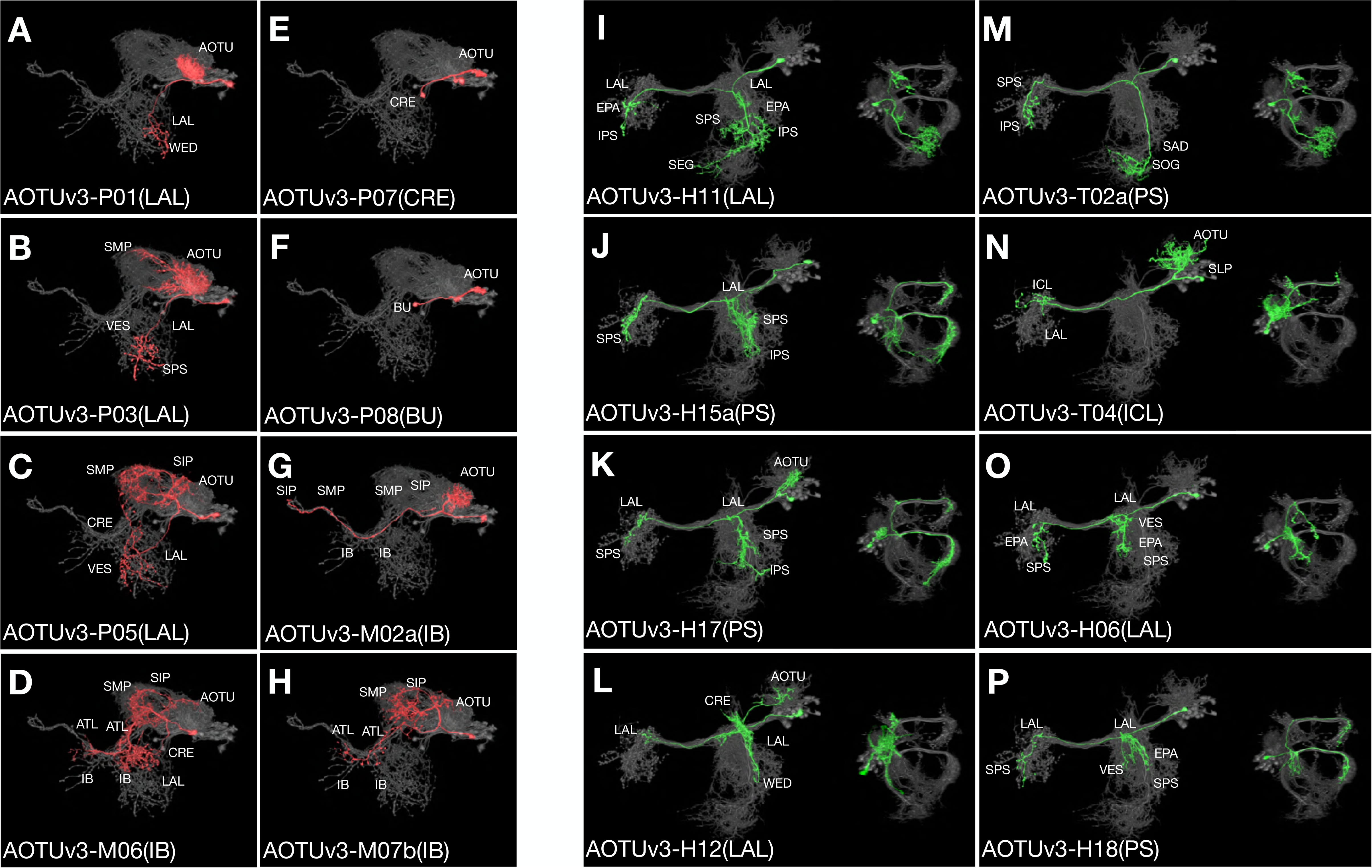
Progressive innervation in AOTUv3B vs. recurrent innervation in AOTUv3A. (A-H) Serially derived single neuron types (red) shown in the context of all recovered AOTUv3B neuron types merged together (grey). Note one M-topology neuron type (D) produced much earlier than many other M-topology types (e.g. [G] & [H]). Both LAL and IB groups show progressive AOTU-to-SP proximal elaborations. (I-P) Serially derived single neuron types (green) shown in the context of all recovered AOTUv3A neuron types merged together (grey). Note discrete extensions at the entry and exit of the shared commissure, and recurrence of similar extension patterns (e.g. [J] & [P]).

By contrast, we see recurrence of characteristic neuron morphology in the AOTUv3A hemilineage (Figure 8I-P). All AOTUv3A neurons project across the brain midline through the same commissure. Their morphological diversity results from variable numbers and paths of major branches that extend out at the common entry and exit points of the commissure. There are three main patterns: 1) two posteriorly projecting branches at both entry and exit points, 2) just one branch at the exit point, or 3) no branch. Various additional features, including ipsilateral AOTU innervation, ipsilateral or bilateral LAL elaboration, and a long ventral extension at the entry point, can exist to increase diversity within each main pattern. Notably, despite midline crossing and largely symmetrical trajectory, there are some T-topology neuron types, connecting ipsilateral SAD with different contralateral neuropils, in this otherwise pure H-topology hemilineage. As to temporal patterning of neuron morphology, we see multiple recurrences of almost all morphological characteristics. For instance, we have recovered neurons with bilateral posteriorly projecting branches in three separate windows, including the first window at the beginning of larval neurogenesis, the second shortly after the first, and the distant third window at the end of the lineage. Nonetheless, certain features (e.g. AOTU innervation) favor early windows, some (e.g. posterior projection only at the exit point) favor late windows, and few (e.g. contralateral ICL targeting) appear in just one window. Taken together, the AOTUv3A hemilineage contains neurons with diverse trajectories in complex temporal patterns.

### Related lineages make analogous neurons in similar temporal patterns

In contrast with drastic differences across paired sister hemilineages, we see striking similarities between certain non-sister hemilineages. Such sharp contrast is well exemplified by comparing the paired AOTUv4A and AOTUv4B hemilineages with the unrelated LALv1A and AOTUv3B hemilineages respectively.

The AOTUv4A hemilineage, like LALv1A, produces only FB-targeting neurons after a short stretch of earlier-born non-FB neurons (Figure 9A-L). Despite targeting only layer 4 to 8 (as opposed to all layers by LALv1A), the AOTUv4A FB neurons resemble the LALv1A FB neurons in both distal and proximal elaborations. We also see similar recurrent targeting of some FB layers in both AOTUv4A and LALv1A. Further, the first AOTUv4A FB neuron type exhibits interesting morphological features characteristic of the last LALv1A FB neuron type. Both show similarly restricted dense elaboration in SP as well as concentrated FB innervation in small subdomains on either the top (AOTUv4A) or bottom (LALv1A) of the FB (Figures 9B and 6L). In conclusion, AOTUv4A and LALv1A make analogous FB neurons with similar or complementary projections in comparable temporal patterns.

**Figure 9.**
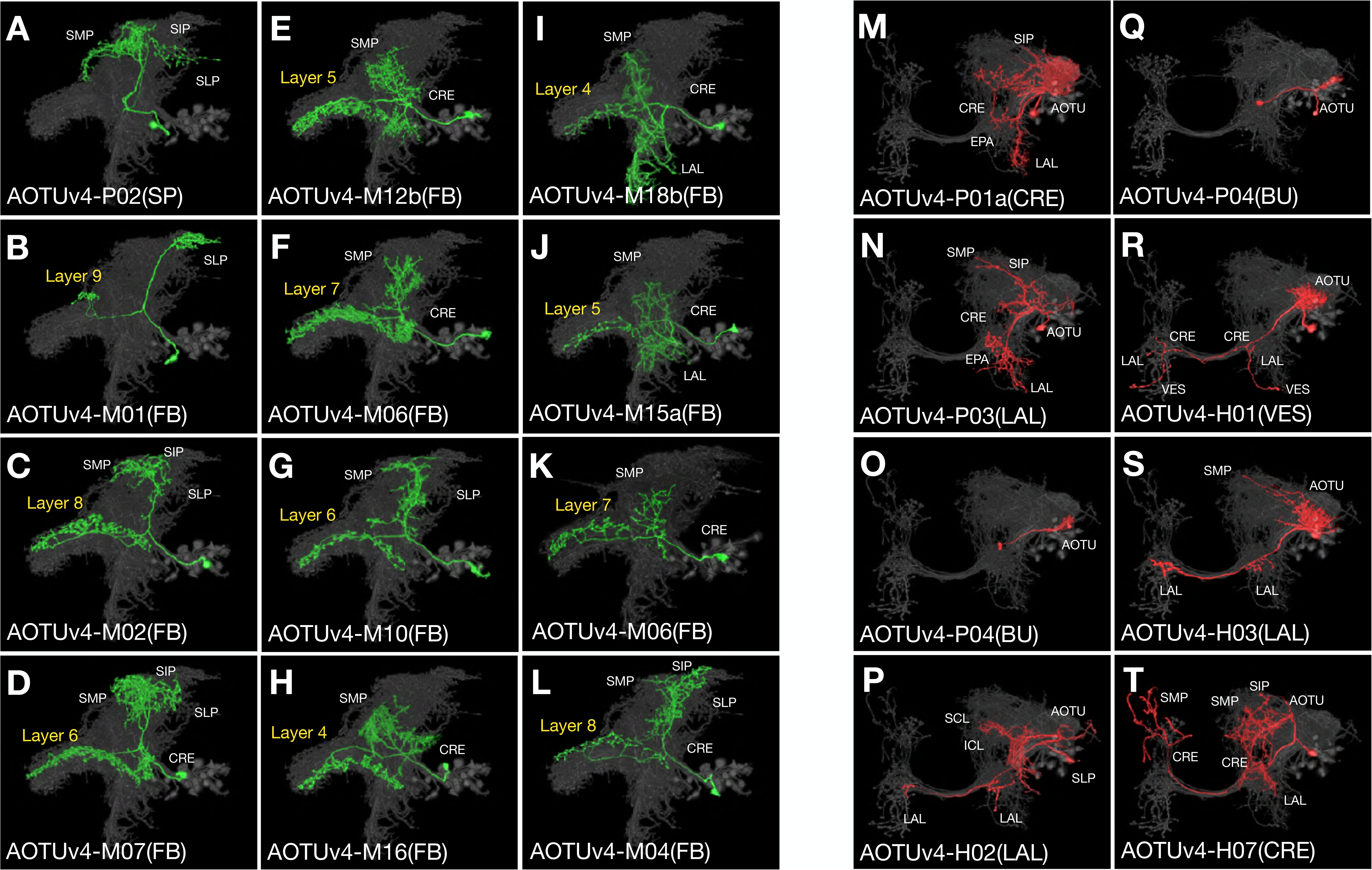
AOTUv4A resembles LALv1A and AOTUv4B resembles AOTUv3B. (A-L) Serially derived single neuron types (green) shown in the context of all recovered AOTUv4A neuron types merged together (grey). Note the early transition from non-CX neurons (A) to FB neurons (B-L), the first FB neuron type innervating layer 9 (B), and recurrent innervation of FB layer 4-8 (C-L). (M-T) Serially derived single neuron types (red) shown in the context of all recovered AOTUv4B neuron types merged together (grey). Note one H-topology neuron type (P) made in the middle of P-topology types (M-O & Q) and much earlier than many other H-topology types (R-T). Both the beginning P-topology CRE group and the later H-topology group show progressive AOTU-to-SP proximal elaborations.

Unlike AOTUv4A, the AOTUv4B hemilineage consists of multiple sequentially produced morphological groups of AOTU-related neurons (Figure 9M-T). This is akin to the AOTUv3B hemilineage. Their similarities exist in spatial as well as temporal patterning of neuron morphology. First, both produce P-topology neurons, then many BU-targeting dot-to-dot neurons, followed by midline-crossing neurons. Second, in the otherwise pure H- or M-topology group of AOTUv4B or AOTUv3B midline-crossing neurons, there consistently exist a few T-topology neurons. Third, we see comparable progressive changes in the proximal elaboration from AOTU to SP in the non-BU groups of both hemilineages. Fourth, in the middle of P-topology neuron production, the appearance of a single neuron type with the H or M topology occurs in both AOTUv4B and AOTUv3B. These extensive parallels between non-sister hemilineages argue for involvement of conserved mechanisms in diversifying neuron fate over time during neurogenesis.

### PAM dopaminergic neurons arise from ‘duplicated’ lineages

In search of closely related lineages, the neighboring CREa1 and CREa2 lineages have long caught our attention because their full-size NB clones show extensive overlapping elaboration in the MB lobes. However, prior to this study, the identities of neurons innervating the MB lobes were elusive. Briefly, we found that each of the NBs produces an intricate sequence of PAM neurons [2] and that one N^on^ hemlineage is indistinguishable from the other.

Both CREa1/CREa2 postembryonic NBs start by making non-PAM N^on^ neurons. Here we ignore the N^off^ neurons, though CREa1B and CREa2B are always distinctive and therefore instrumental to distinguishing CREa1A from CREa2A. The first CREa1A neuron connects the ipsilateral CRE/LAL with the contralateral SMP/ATL/IB. By contrast, the first CREa2A neuron dies prematurely, leaving its identity unclear. However, the next two surviving neuron types in CREa1A and CREa2A both innervate the FB. Strikingly, the CREa1A FB neurons are morphologically indistinguishable from the CREa2A FB neurons. We could tell them apart only by determining their paired sister neurons.

Following the brief production of morphologically identical FB neuron types, both CREa1A and CREa2A NBs yield PAM neurons until they exit the cell cycle. Despite the serial production of different PAM neurons, the CREa1A and CREa2A PAM neurons (identified based on their paired sister neurons) look identical at all time points. This phenomenon indicates that the two NBs produce two identical series of PAM neurons.

In each series of PAM neurons, we identified 17 types of PAM neurons based on MB lobe innervation patterns. This list covers all 14 previously reported types of PAM neurons plus three undocumented morphological types (r4<r2, r4r5, and b’2r5). In addition, we observe variants of b2, r5, r4, and b’2p that arise in separate windows and show birth time-dependent patterns of dendrite elaboration (Figure 10). By analyzing birth order, we see progressive innervation of neighboring zones in the MB medial lobes. a1 is innervated first and the pattern progresses medially along the beta lobe, then backwards while alternately innervating the gamma and beta’ lobes, then medially again along the gamma lobe, and finally ending in b’2 and b2 (Figure 10A). In conclusion, the PAM cluster of dopaminergic neurons arise in duplicated series from the non-sister CREa1A and CREa2A hemilineages.

**Figure 10.**
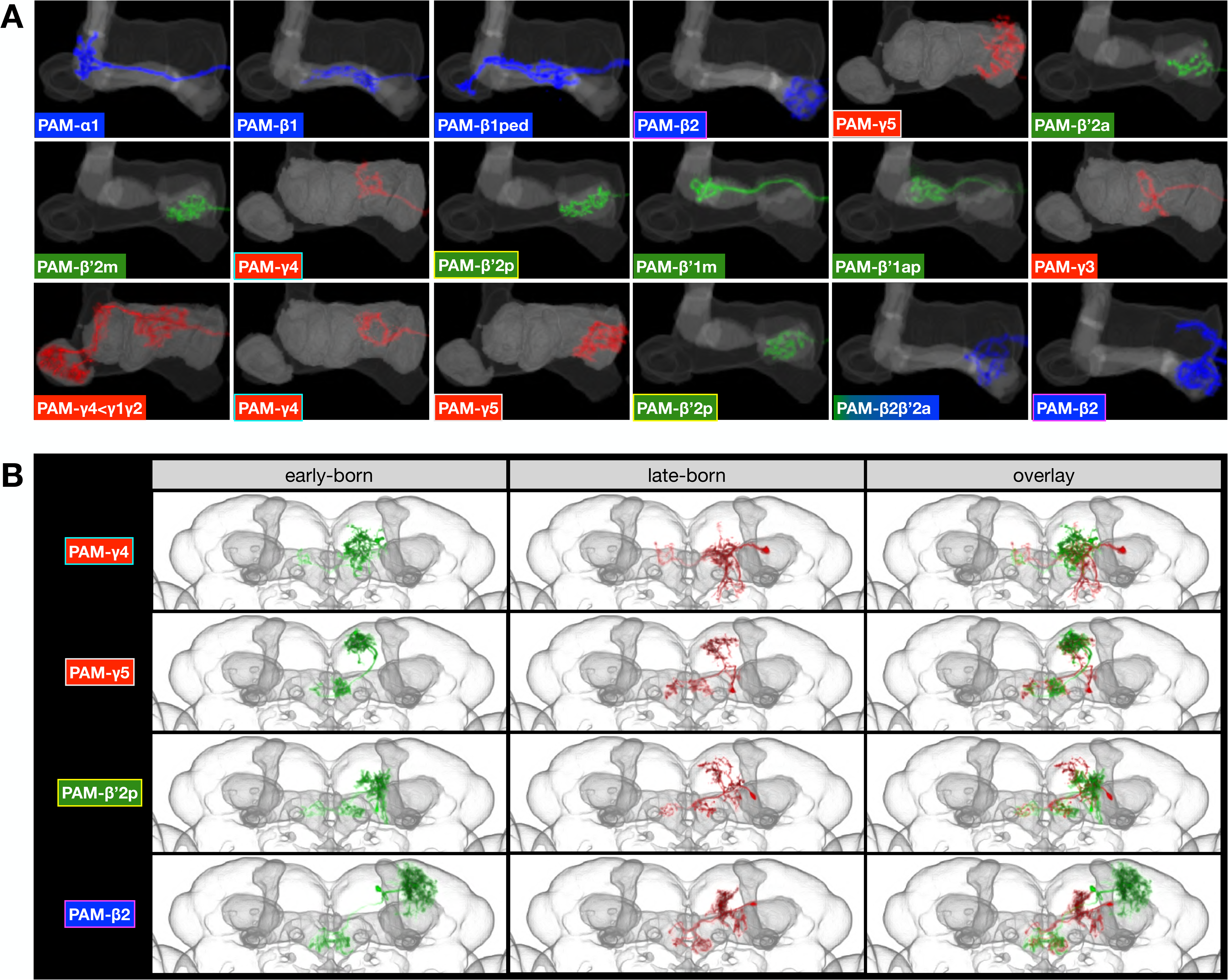
Patterned innervation of MB medial lobes by serially derived PAM neurons in both CREa1A and CREa2A. (A) PAM neuronal elaborations (pseudo-colored based on lobe identity) within the MB medial lobes (grey) showing progressive innervation of neighboring MB lobe zones. Note recurrence of four targeting patterns: beta2, gamma5, gamma4, beta’2p. (B) Representative PAM neurons (green-colored ones born before those in red) merged onto the same adult brain template. Note birth time-dependent proximal elaborations in neurons targeting the same zone.

## Discussion

A series of fating events occur to promote diverse neuronal types. First, NBs acquire lineage identity via spatial patterning cues, then sequentially born GMCs inherit temporal factors, and finally GMCs divide into pairs of distinct neurons, generating sister hemilineages with differential Notch signaling. Predetermined fates evidently guide all aspects of neuronal differentiation in the invariant fly lineages [23]. Despite having predetermined fates, it is hard to imagine how complex neuronal morphology is controlled. Mapping neuron morphology for 25 hemilineages in this study reveals that primary trajectories and thus neuropil targets are dependent upon both lineage identity and Notch signaling (*i.e.* hemilineage identity). Neuronal temporal patterning, by contrast, governs the sequence of targeting various neuropils in those hemilineages with multiple targets. Temporal fates further diversify neuronal elaborations within the target in neuropil-characteristic manners.

The nature of neuronal temporal fate specification appears conserved across distinct hemilineages. This is evidenced by the presence of common temporal features in diverse hemilineages. For instance, many hemilineages possess first-born neurons with unique extended projections and/or last-born neurons with simplified morphology. Additional general temporal patterning phenomena exist. For instance, most hemilineages comprised of multiple morphological neuronal classes alter neuropil targeting in a progressive manner. Also, we see cyclic morphological changes or the recurrence of characteristic morphology in many hemilineages, especially those involved in building fine topographic maps. Furthermore, unrelated hemilineages can be remarkably similar, even though the sister hemilineages are clearly divergent. This is best exemplified by the respective analogies between the AOTUv4A and LALv1A hemilineages and the AOTUv4B and AOTUv3B hemilineages. Notably, the FB neurons of AOTUv4A and LALv1A show similar, but not identical, patterns of neurite elaboration. This suggests that terminal arborization can also be directly or indirectly regulated by hemilineage identity. Nonetheless, their temporal complexities are comparable. Taken together, our data suggest that hemilineage identity governs differential manifestation of temporal fates primarily through regulating neuropil targeting.

As to possible molecular mechanisms, we speculate that hemilineage identity regulates expression of specific targeting receptors which are further subject to the control of temporal factors. For instance, the known temporal gradients of Imp, Syp, and Chinmo [24] [25] may underlie differential expression of hemilineage-dependent axon guidance receptors over time, to achieve progressive targeting of distinct neuropils by serially derived neurons. About intra-neuropil patterning, the temporal diversification of terminal elaboration is neuropil-specific and largely hemilineage-independent. This suggests involvement of universal targets downstream of common temporal factors that can versatilely support a wide spectrum of neuropil-characteristic morphological diversity. In addition, the known temporal factors Cas and Svp, which are expressed in NBs during early postembryonic neurogenesis [26], could promote the extended projections of first-born neurons. The cyclic/recurrent temporal fates may result from dynamic Notch signaling, reminiscent of Notch-dependent alternate production of AL and AMMC projection neurons in the lateral AL lineage [19]. In sum, there likely exist conserved temporal fating mechanisms that play a central role in lineage-guided neuronal morphogenesis.

Given the need for distinct hemilineages in making diverse sets of related neuron types, it is interesting to see the production of two long identical series of dopaminergic neuron types by CREa1A and CREa2A. Strikingly, their homology extends throughout postembryonic neurogenesis. The two FB neuron types born prior to dopaminergic neurons in CREa1A are also morphologically indistinguishable from those in CREa2A. The only difference is the programmed death of the first larval-born CREa2A neuron. In contrast, their paired sister hemilineages (CREa1B and CREa2B) are easily distinguishable as all and only CREa1B neurons cross the midline. These phenomena implicate that neighboring CREa1 and CREa2 lineages may have arisen from NB duplication followed by a change in midline crossing. Thus, we believe that one way for brain complexity to increase is through lineage duplication and subsequent divergence.

In conclusion, mapping parallel neuronal lineages with single-cell resolution in intact brains reveals how a complex brain can be reliably built from differentially fated neural stem cells. This seminal groundwork lays an essential foundation for unraveling brain development from genome to connectome.

## Materials and Methods

### Fly strains and DNA constructs

Transgenes used for twin-spot MARCM for vnd lineages include: vnd-T2A-GAL4 (this study), UAS-KD, dpn>KDRT-stop-KDRT>Cre:PEST, nSyb>loxP-stop-loxP>LexA::p65, hs-FLP, FRT40A, lexAop-mCD8::GFP-insulated spacer-lexAop-rCD2i, and lexAop-rCD2::RFP-insulated spacer-lexAop-GFPi [16]. Transgenes used for Notch depletion include: hs-ATG>KOT>FLP [27], ase-KD (this study), dpn>FRT-stop-FRT>Cre::PEST [16], actin^loxP-stop-loxP^Gal4 (this study), UAS-Notch-RNAi (BL#33611), USA-mCD8::GFP [10].

vnd-T2A-Gal4, homology arms of about 3 kb each were cloned into pTL1 for knocking-in T2A-Gal4 in vnd with following primers: vnd_55AgeI: TACGACCGGTGATCAAGGAGAACGAGCTATACG; vnd_53StuI: AAGGCCTGGGCCACCAGGCGG; vnd_35PmeI: TACGGTTTAAACTAATATTGCTAGGAACTGGCATTCAC; vnd_33MluI: AAGTACGCGTAACTGGAATAAGTTC. T2A-Gal4 CDS was inserted right after the second last amino acid using traditional Golic heat shock strategy for gene targeting to obtain vnd-T2A-Gal4 transgenic fly [28].

ase-KD, the asense promoter [29] was put in front of the KD in modified pBPGw through gateway system (Invitrogen) as described previously [16].

actin^loxP-stop-loxP^Gal4, a synthetic Flox cassette was inserted into a KpnI site between the *actin* promoter and Gal4 as described previously [16].

### MARCM

For ts-MARCM clonal analysis, 0–2 h old newly hatched larvae with proper genotype were collected and put into vials (80 larvae/vial) containing standard fly food. The larvae were raised at 25°C until desired stages. Organisms were resynchronized with respect to puparium formation for those clones induced at late larval and early pupal stages. To induce clones, the organisms were heat-shocked at 37°C for 15–40 min. After heat shock, the organisms were put back to 25°C until dissection. For Notch depletion, newly hatched larvae with proper genotype were heat shocked at 37°C for 15 min to induce the activation of lineage restricted driver for clonal labeling and Notch depletion. After heat shock, the larvae were put back to 25°C until dissection.

### Immunostaining and Confocal Imaging

Adult brains were dissected, fixed, and processed as described previously [16]. Antibodies used in this study include rabbit anti-GFP (1:1,000, Invitrogen), rat monoclonal anti-mCD8 (1:100, Invitrogen), rabbit anti-RFP (1:1,000, Clontech), mouse monoclonal anti-Bruchpilot, nc82 (1:50, Developmental Studies Hybridoma Bank), Alexa 488, (Invitrogen), Cy3, Cy5 or Alexa 647 (Jackson ImmunoResearch) conjugated anti-mouse, anti-rabbit, and anti-rat antibody (1:500). After immunohistochemistry, brains were post-fixed with 4% PFA in PBS for 4 hr at RT followed by four washes in PBT for 10 mins and then rinsed with PBS. Brain samples were placed on poly-L-lysine-coated cover slips followed by series dehydrated in ethanol baths (30%, 50%, 75%, 95%, and 3 × 100%) for 10 min each and then 100% xylene three times for 5 min each in Coplin jars. Samples were embedded in DPX mounting medium (Electron Microscopy Sciences, Hatfield, PA). Fluorescent signals of whole-mount adult fly brains were acquired at 1 μm intervals using a 40× C-Apochromat water objective (NA=1.2) and 0.7 zoom factor at 1024×1024 pixel resolution on Zeiss LSM710 confocal microscope (Carl Zeiss). The “Janelia Workstation” image-viewing software (Murphy et al., unpublished data) was used to analyze confocal stacks. The whole brain images were registration and alignment to a standard brain (JFRC2013) using the reference nc82 channel as described previously [2].

## Acknowledgements

We thank Janelia Workstation, Janelia FlyLight, and Janelia Fly Core for technical supports. We thank Masayoshi Ito and Jens Goldammer for comments. We thank Crystal Di Pietro and Kathryn Miller for administrative support. This work was supported by Howard Hughes Medical Institute.

